# Conservation genomics reveals low genetic diversity and multiple parentage in the threatened freshwater mussel, *Margaritifera hembeli*

**DOI:** 10.1101/2020.03.31.018689

**Authors:** Nicole L. Garrison, Paul D. Johnson, Nathan V. Whelan

## Abstract

*Margaritifera hembeli* is a federally threatened freshwater mussel species restricted to three central Louisiana drainages. Currently, management efforts are being formulated without an understanding of population-level genetic patterns, which could result in sub-optimal conservation outcomes. In particular, information about riverscape genetic patterns is needed to design effective propagation and reintroduction plans. We apply a genomic approach (RADseq) to assess genetic diversity and structure among four wild populations sampled from across the species range. We also assess the genetic diversity of a captively reared cohort produced from a single female. We recovered population differentiation between individuals sampled to the north and south of the Red River. All sites had similarly low levels of heterogeneity and other measures of genetic diversity. The captive cohort displayed higher levels of genetic diversity than expected and likely represents a case of multiple paternity. Future propagation efforts will likely be able to produce genetically diverse cohorts from a small number of wild-caught females, and we recommend future reintroduction efforts utilize brooders within the sub-drainage closest to the reintroduction effort.

## Introduction

In a world of ever-increasing anthropogenic encroachment and climatic change, effective conservation decisions for species of concern must be made quickly. Regardless of taxon, detailed genetic assessments clarify the pattern and process of diversity across a landscape, diagnose specific conservation challenges, and answer crucial questions managers face during the decision making process (DeSalle and Amato 2004; Segelbacher et al. 2010; Richardson et al. 2016). Genetic information is invaluable to captive propagation programs, where data play a role in the selection of broodstock and maintenance of genetic diversity over time (Witzenberger and Hochkirch 2011). In cases where propagation methods are actively being developed and reintroduction efforts planned, genetic analysis can assist production and brood stock selection, better insure successful reintroduction outcomes, and enable continued monitoring of established populations (Jones et al. 2006; Schwartz et al. 2007; McMurray and Roe 2017). Despite the potential contribution of genetic information to successful conservation efforts, many species have little or no genetic data available; this is particularly true of freshwater mussels(Haag and Williams 2014; Strayer et al. 2019).

Conservation genetic information can now be inferred using thousands of markers from across a genome, providing a high resolution alternative to historic standbys like single gene or microsatellite analyses (Luikart et al. 2003; Davey and Blaxter 2010). Importantly, high throughput methods can be employed in non-model systems with no pre-existing genomic resources. Population genomic analyses can also directly aid in the development of markers for continued monitoring of wild and captive populations (Schwartz et al. 2007; Karlsson et al. 2011; Hendricks et al. 2018) and they represent a powerful tool for conservation managers and stakeholders overseeing long-term recovery programs (Witzenberger and Hochkirch 2011). Leveraging the power and accessibility of current conservation genomic methods to examine understudied groups such as freshwater mollusks should be a major focus of conservation research as anthropogenic pressures increase.

Freshwater mussels are among the most critically imperiled aquatic organisms. A history of overexploitation and environmental impacts including habitat fragmentation, channel alterations, agricultural inputs, and pollutants has led to global endangerment of mussels. In the last 100 years, an estimated 28 species have gone extinct in North America alone. Of the remaining North American species, approximately 65% are currently at risk of extinction (Williams et al. 1993; Haag and Williams 2014). The importance of mussels to freshwater ecosystems globally cannot be overstated. Mussels are long-lived benthic invertebrates that actively filter suspended particles from water, providing valuable ecosystem services (Vaughn and Hakenkamp 2001). Furthermore, freshwater mussels are intimately linked to water quality, and their shells can provide a record of environmental changes, at both recent (Pfister et al. 2011) and historic scales (Fritts et al. 2017). As part of their complex life cycle, mussels produce parasitic larvae (glochidia) that must attach and mature on the gills of a host fish. This relationship can sometimes be very specific (i.e. a single host species) or more general (a particular family of fishes), and is required for the maturation and dispersal of juvenile mussels (Wächtler et al. 2001). A history of overexploitation and environmental impacts including habitat fragmentation, channel alteration, agricultural inputs, and pollution has led to global endangerment of mussels (Lopes-Lima et al. 2018b). Environmental degradation undoubtedly impacts mussels directly through physiological stress (Strayer et al. 2004), but may also impair recruitment via the reduction of host fish population densities (Bogan 1993; Haag 2012) and their ability to migrate freely (Watters 1996).

*Margaritifera hembeli*, Louisiana Pearlshell (Fig. 1a), is federally threatened under the U.S. Endangered Species Act and is restricted to three tributary drainages of the Red River in central Louisiana (Smith 1988)(Fig. 2). Belonging to the family Margaritiferidae, *M. hembeli* is a morphologically and phylogenetically distinct group of mussels with only five extant North American representatives, much fewer than the number of mussel species representing the family Unionidae (Bogan 2008; Lopes-Lima et al. 2018a). *Margaritifera hembeli* individuals are long lived (~45-75 years) and inhabit shallow, high velocity stream reaches with relatively stable substrates (Johnson and Brown 1998; Johnson and Brown 2000). Healthy occurrences of this species are characterized by large, dense beds (Fig. 1b), sometimes exceeding 300 individuals/m^2^ (Johnson and Brown 1998). In the wild, *M. hembeli* glochidia have been found on *Noturus phaeus* (Brown Madtom), *Luxilus chrysocephalus* (Striped Shiner), *Lythrurus umbratilis* (Redfin Shiner), and *Notemigonus crysoleucas* (Golden Shiner), but these may be spurious records and overall host suitability has not been confirmed (Hill 1986; Johnson and Brown 1998). In particular, host fish specificity in the wild remains unclear given that *M. hembeli* glochidia have only been found in small numbers on implicated fish species, yet they transform particularly well on *Esox* spp. species in captivity (see below). Many negative factors influencing mussels on a global scale are also affecting *M. hembeli*. The 1988 United States Fish and Wildlife (USFWS) status report for the species noted that *M. hembeli* populations face threats from poor land management practices (silviculture, gravel mining), reservoir construction, and pollution runoff. During this initial USFWS assessment of *M. hembeli*, it was known only from 11 headwater streams in Bayou Boeuf (Rapides Parish, Louisiana), which contributed to the designation of endangered status under the U.S. Endangered Species Act (USFWS 1988). Upon the discovery of additional populations in Grant Parish, it was later down listed to threatened (USFWS 1993). Nevertheless, *M. hembeli* remains in urgent need of conservation attention.

**Fig. 1.**
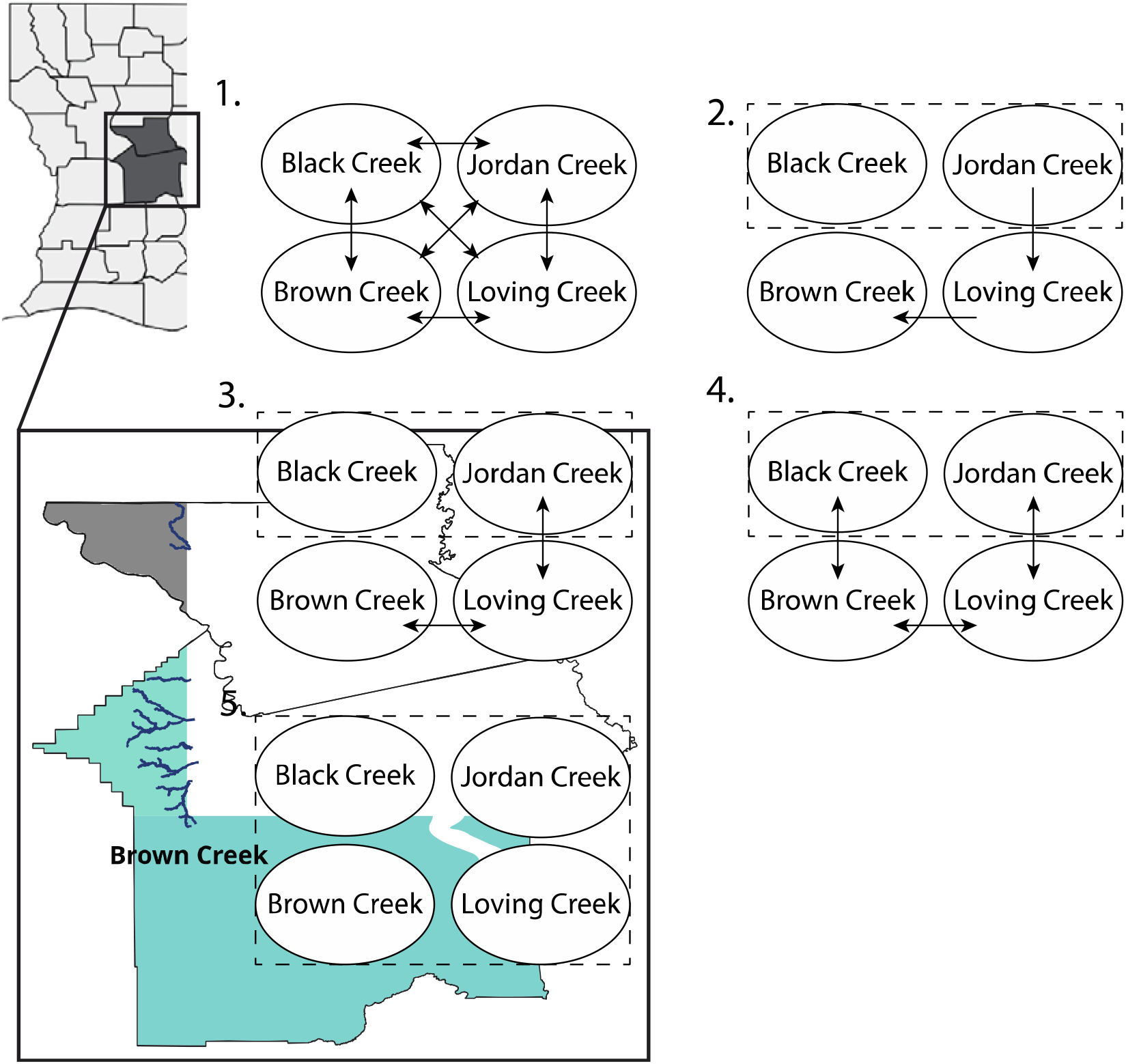
Map depicting the extent of M. hembeli distribution in central Louisiana. Pictures illustrate mussel bed density at Black Creek site (upper) and individual shell (lower).

**Fig. 2.** Population migration models assessed using migrate-N.

For *M. hembeli* and other mussels, there is considerable interest in captive propagation as a management tool (Haag and Williams 2014). Currently, most mussel propagation programs involve the capture of one or a few wild, gravid females followed by inoculation of glochidia on the appropriate host fish in captivity. Progress has been made in technical aspects of rearing certain species such as determining temperature thresholds (Steingraeber et al. 2007), appropriate nutrition (Gatenby et al. 1996), sediment requirements (Jones et al. 2005), host fish identification (Keller and Ruessler 1997; Hart et al. 2018), and creating capacity for large scale production (Barnhart 2006). However, genetic impacts of propagation are less clear. Though many propagation operations exist at federal and state government facilities across the southeast, genetic consequences of in-use propagation protocols for freshwater mussels is an area of research in its infancy. Recommendations exist (Jones et al. 2006), but few studies have explicitly examined the genetic consequences of specific propagation protocols for freshwater mussels.

An initial transformation and culture trial for *M. hembeli* producing individuals utilized in this study began in 2016. Transformation of *M. hembeli* juveniles was completed on a single *Esox americanus americanus* (Redfin Pickerel) by USFWS (Natchitoches National Fish Hatchery in Louisiana and the Ecological Services Office in Louisiana). Infection rate was robust and the effort resulted in nearly 9,000 juvenile *M. hembeli* transformed from the single *E. americanus americanus*. Once transformation had been completed, culture protocols were initiated at the Alabama Aquatic Biodiversity Center (AABC) in Marion, Alabama to evaluate three different methods for rearing *M. hembeli* juveniles. As this initial effort was focused primarily on determining basic culture methods for *M. hembeli*, transformation of juveniles from a single gravid *M. hembeli* from Black Creek (Grant Parish, Louisiana) represented an opportunity to examine genetic variation of progeny found in a single female. Past research has indicated females of other *Margaritifera* species can mate with multiple males in the wild (Wacker et al. 2018) and multiple paternity has been noted in several Unionid mussel species (Christian et al. 2007; Bai et al. 2012; Ferguson et al. 2013). No study has examined this potential mating system in *M. hembeli*. Given conservation challenges facing *M. hembeli*, the prospect of reintroducing large numbers of low diversity individuals should be carefully considered. Understanding of both the genetic structure of wild populations and the opportunity to contrast against the diversity of a single-female brood could prove invaluable for informing future conservation and management recommendations for *M. hembeli* reintroduction efforts.

Despite genetic concerns for the management and survival of *M. hemebli,* modern genetic data has not been generated to inform management decisions. Previous genetic work on *M. hembeli* found extreme monomorphism (H_o_ = 0) across 25 allozyme markers (Curole et al. 2004). This observed near-absence of heterozygosity was hypothesized to result from the common (and likely natural) stochastic extirpation of *M. hembeli* beds, coupled with high re-colonization rates. A more recent study utilizing microsatellites (Roe 2009) found low heterozygosity at all sampled populations and some genetic structure among *M. hembeli* populations, but conclusions were drawn from only five loci. In total, previously generated genetic data offer scant information for use in designing management efforts. Modern, genomic-level data are necessary to assess population connectivity, measure genome-wide genetic diversity, and provide managers with useful information that can be used to inform propagation and reintroduction efforts.

The goals of this study were to 1) assess the genetic diversity of the 2016 AABC captive *M. hembeli* cohort 2) determine whether previous reports of low genetic diversity of wild *M. hembeli* populations using allozymes is supported by genomic markers and 3) and characterize patterns of gene flow or potential barriers to aid in reintroduction of future culture efforts. To accomplish these objectives, a restriction enzyme associated high-throughput sequencing method was employed (RADseq) to generate a dataset of thousands genomic loci. RADseq is capable of utilizing genomic DNA samples collected non-lethally to generate millions of sequence reads for each individual. These reads can be processed with existing tools to identify thousands of single nucleotide polymorphisms (SNPs) from across the genome, providing high resolution data for determining demographic and evolutionary dynamics within a species. Ultimately, these data will inform recovery efforts of *M. hembeli* and provide the first genetic profile of a propagated cohort for this threatened freshwater mussel.

## Materials and Methods

### Sample Collection and Sequencing

We sampled 20 individuals each from four sites in the Red River drainage in Louisiana, and 20 captively reared individuals produced from a single wild caught female from Black Creek. Two sampling localities were from Rapides Parish (Brown Creek – Bayou Rapides and Loving Creek – Bayou Boeuf) and two were from Grant Parish (Jordan Creek and Black Creek – Bayou Rigolette). Sampling sites were selected for ease of access, abundance of *M. hembeli*, and because they encompass the major drainages from which *M. hembeli* is known to occur (Fig. 2). Mussels were collected by hand and effort was taken to select a broad size range likely representing individuals from multiple age classes. A sterile, individually wrapped buccal swab was used to collect genetic material from the foot of each mussel and immediately placed in swab stabilization buffer from the Buccal-PrepPlus DNA isolation kit (Isohelix^™^). Mussels were photographed on site and returned immediately to their bed after swabbing. The captively reared individuals were lethally sampled at the AABC and a clip of the foot was placed in 96-100% ethanol until DNA extractions could be completed. The mother of the captive cohort was not available for sampling and a tissue voucher/buccal swab was never taken. Shells of captively reared individuals were deposited at the Auburn Museum of Natural History (AUM 45578-45595).

DNA extractions of buccal swabs and tissue were completed with Isohelix Xtreme DNA isolation kit and the Qiagen DNeasy blood and tissue kit respectively, following the manufacturers’ instructions. After DNA extraction, each sample was treated with RNase A at a final concentration of 100 μg/ml and incubated at 37°C for 15 minutes to remove any co-purified RNA. Each extraction was quantified with a Qubit Fluorometer and checked for integrity of high molecular weight DNA through standard gel electrophoresis. Samples were standardized to a concentration of 20 ng/μL and 50 μL of standardized DNA was sent to Floragenex Inc. (Portland, OR) for RADseq library preparation using the SbfI restriction enzyme following Baird et al. (2008). Samples were tagged with unique barcode identifiers, pooled, and sequenced in three replicate lanes on the Illumina HiSeq 4000 platform using 100bp paired-end chemistry.

### Sequence Processing and Variant Identification

Reads from all three sequenced lanes were combined for each individual and processed with the STACKS v2.1 pipeline for population genomic analysis (Rochette et al. 2019). STACKS demultiplexes raw sequencing data, aligns reads to form stacks of loci, identifies variants (i.e. SNPs), and generates descriptive population statistics. Paired-end reads were demultiplexed with the *process_radtags* command using default settings that allow for barcode sequences to be rescued if the barcode varies by only one nucleotide from the expected sequence. Input file preparation details and *process_radtags* settings can be found online (https://github.com/nlgarrison/ConservationGenomics). The denovo_map.pl script was used to automate the STACKS pipeline, as a reference genome is not available for *M. hembeli* or any closely related species. Stack assembly required a minimum of five reads per locus (-m 5), allowed for three mismatches within stacks of the same individual initially (-M 3) and two mismatches between stacks of different individuals (-r 2). All other parameters were set to default. To identify SNPs in stacked loci, a catalog of potential sites must be formed; in our pipeline only individuals from the wild populations were used to generate the catalog. To do this, at the *cstacks* step in the denovo_map.pl pipeline, a population map including only wild-caught individuals was provided to the catalog building command. This approach was chosen to reduce bias in called variants that might be introduced by uneven population sampling, and it allowed for a true test of population assignment for the captive individuals.

We ran the *populations* command on two sets of samples: wild-caught individuals only (NoCaptive) and all sampled individuals (WithCaptive). The captive cohort was excluded in initial runs of *populations* to eliminate bias when estimating the baseline population parameters for wild individuals. The NoCaptive dataset was analyzed with a minimum minor allele frequency setting of 0.025, maximum heterozygosity setting of 0.50, and a requirement that a variant be present in at least three sampling sites and at least 50% of individuals within each sampling site; these settings were also used in the analysis of the WithCaptive dataset. For both the NoCaptive and WithCaptive datasets, one *populations* analysis was generated allowing only one random SNP per locus (denoted “S”) and a second was produced allowing multiple SNPs per locus (denoted “M”), resulting in a total of four datasets; NoCaptiveS, NoCaptiveM, WithCaptiveS, WithCaptiveM. This was done because some downstream analyses assume unlinked loci, whereas others can use multiple SNPs originating from the same locus. For downstream analyses requiring subsets of individuals as input, the program VCFtools (Danecek et al. 2011) was used to generate reduced datasets as needed from the WithCaptiveS/M vcf files.

### Population Genomic Analyses

For each dataset, average heterozygosity, nucleotide diversity, pairwise F_ST_ among each sampling site, and F_IS_ at each sampling site was reported by *populations*. The *basicStats* function of the R (R CORE Team 2019) package diveRsity (Keenan et al. 2016) was used to calculate allelic richness. We assessed population structure among sampling sites with an Analysis of Molecular Variance (AMOVA; Excoffier, Smouse, and Quattro 1992) and a series of clustering methods. AMOVA was performed using the *poppr.amova* command in the R package adegenet (Kamvar et al. 2014; Jombart et al. 2018). Individuals were stratified by sample site and whether the site was located north or south of the Red River to assess whether the Red River serves as a barrier to gene flow or if genetic structure is better explained by local landscape characteristics. Significance was tested with a 500 permutation randomization test.

Discriminant analysis of principal components (DAPC) was implemented using the R package adegenet following Jombart and Collins (2015), using the NoCaptiveM and WithCaptiveM datasets. The best-fit number of clusters (K) was assessed using *k*-means clustering with Bayesian information criteria. The *snmf* function of the R package LEA (Frichot and François 2015) was used to identify clusters of individuals and determine admixture proportions, and the best-fit K was determined using the cross-entropy criterion. For these analyses, only the unlinked SNPs (NoCaptiveS, WithCaptiveS) were used. Though similar in function to STRUCTURE (Pritchard et al. 2000), LEA can be more accurate than STRUCTURE in the face of inbreeding (Frichot et al. 2014) and handles genomic data more efficiently.

We also used the model-based method fineRADstructure (Malinsky et al. 2018) to generate a summary of haplotype coancestry for all individuals. A major advancement provided by fineRADstructure is its ability to utilize linkage and polymorphism in genomic data without a reference genome, allowing fine scale patterns of relatedness among individuals to be examined. This analysis was conducted with the NoCaptiveM, WithCaptiveM, and the captive cohort in isolation. The captive cohort was examined independently in order to make more accurate inferences about potential multiple paternity.

We tested for a signature of isolation-by-distance using a Mantel test of correlation between geographical distance and pairwise F_ST_ values for the wild populations. Geographic distances among sites were measured by plotting sample collection points in QGIS (QGIS Development Team 2014) and hand tracing stream distance between sampling sites in a pairwise fashion. As river connections were sometimes difficult to assess and to account for minor idiosyncrasies associated with tracing river path, hand tracing was repeated three times for each pair and an average of distances in meters was taken as the geographic stream distance. F_ST_ values used for the Mantel test were those reported by STACKS (Table 1). The Mantel test was done using the R package ‘ade4’ (Dray, Dufour, et. al 2007); significance was evaluated with 1,000 random permutations. As Mantel tests have received criticism for use as a measure of isolation by distance (Legendre et al. 2015; Meirmans 2015), we also performed a multiple regression on distance matrices with the MRM function of the Ecodist R package (Goslee and Urban 2007); significance was tested with 10,000 permutations.

**Table 1.** Pairwise FST (top) and estimated in-stream distance in km (bottom) for all populations analyzed.

|  | **Jordan** | **Black** | **Loving** | **Brown** | **Captive** |
| --- | --- | --- | --- | --- | --- |
| **Jordan** | - | 0.043 | 0.063 | 0.078 | 0.048 |
| **Black** | 36.5 | - | 0.064 | 0.082 | 0.035 |
| **Loving** | 77.1 | 91.4 | - | 0.063 | 0.071 |
| **Brown** | 87.8 | 102.5 | 41.2 | - | 0.086 |
| **Captive** | 36.5 | 0 | 91.4 | 102.5 | - |

The programs diveRsity and Migrate-n version 4.2.14 (Beerli and Palczewski 2010) were used to evaluate connectivity between wild populations of *M. hembeli*. The *divMigrate* function of the R package diveRsity uses differences in allele frequencies to model asymmetric, relative rates of migration between populations (Sundqvist et al. 2016). Though it was designed for use with microsatellite data, *divMigrate* is capable of handling genomic SNP datasets and has been shown to reflect biologically realistic scenarios of population connectivity in recent studies (e.g. Woodings et al. 2018; Manuzzi et al. 2019). An analysis of the WithCaptiveS dataset was done with the *divMigrateOnline* implementation (Keenan 2012) to calculate relative rates of migration between wild populations using a cutoff value of 40, an alpha of 0.05, and the G_ST_ statistic (Nei 1973). Support for the asymmetry of migration rates was evaluated with 1000 bootstrap replicates. Although the captive population was included in our initial *divMigrate* analysis, due to its high similarity (G_ST_ = 1.0) with Black Creek individuals it was masked using the “exclude population” option without recalibrating rates.

The *divMigrateOnline* analysis was complemented by the Bayesian population genetics program migrate-n. Given computational demands of migrate-n, a subset of 100 polymorphic loci were randomly selected from the NoCaptiveM dataset. Migrate-n analyses were done with loci in their entirety, rather than individual SNPs, as the SNP model has not been thoroughly tested (see migrate-n manual). Loci that appeared more than once in the random subset were filtered out, leaving 95 polymorphic loci for inclusion in the migrate-n analysis. Geographic distances between sites were calculated as described previously. Five migration models were investigated as indicated by geography and other population genomic analyses 1) full migration 2) northern panmixia with unidirectional gene flow from Loving Creek to Brown Creek 3) northern panmixia with bidirectional gene flow between Loving Creek and Brown Creek 4) panmixia (Fig. 3). For each migration model, Bayesian inference was performed using the DNA sequence model (sampling of 20,000,000 total steps, 10,000 steps discarded as burn-in, default priors) and combined with thermodynamic integration (four parallel heated chains) at temperatures 1, 1.5, 3.0, and 1×10^6^. Log marginal likelihood values were calculated with Bezier approximation within migrate-n and log Bayes factors were used to rank the models following Beerli & Palczewski (2010).

**Fig. 3.**
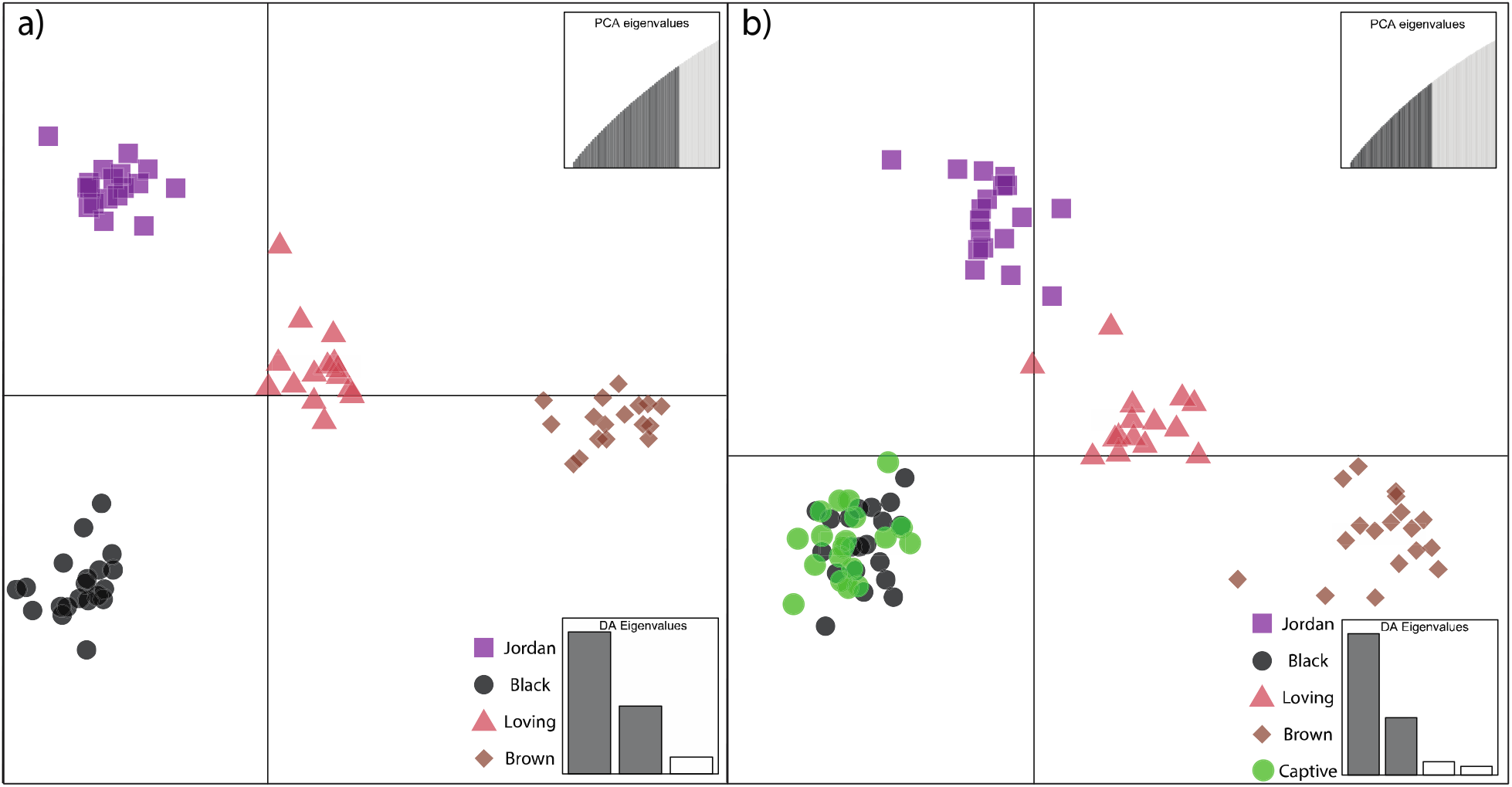
Discriminant analysis of principal components output showing clustering of a) all wild populations and b) wild + captive individuals using the multiple SNP per locus datasets.

Effective population size (N_e_) and the number of effective breeders (N_b_) was estimated for each sampling site, geographically proximate wild populations combined, and the captive cohort in isolation with NeEstimator2 (Do et al. 2014) using the NoCaptiveS and WithCaptiveS datasets as input. Given results indicating that the Red River represents a barrier to gene flow (see below), we combined sites to assess N_e_ of “northern” (Black Creek and Jordan Creek) and “southern” (Loving Creek and Brown Creek) populations. The molecular coanscestry method of Nomura (2008) and linkage disequilibrium method (Hill 1981; Waples 2006) as implemented in NeEstimator2 were used to evaluate N_e_ and N_b_ without a genomic map.

The program COLONY (Jones and Wang 2010) was used to validate suspected multiple paternity in the captive cohort. A captive-only subset of SNPs filtered from the WithCaptiveM dataset using vcftools was used as input. Three different filtering strategies were tested, allowing sites with missing data proportions of 0%, 25%, and 50%. Files were converted to COLONY input format using tidy_vcf and the write_colony function of the R package radiator (Gosselin et al. 2020). Long, full likelihood runs were performed with 10 replications for each filtered dataset; male and female polygamy settings were used and half sibship of all individuals with an unknown mother was specified in the input file.

## Results

Sequencing resulted in >460 million reads per lane and >1 million reads per individual (n=94). Sequence reads filtered out due to barcode ambiguity, unclear restriction cut sites, and potential Illumina adapter contamination comprised ~15% of the total raw reads; 1.4 billion paired-end reads were included in the assembly process. We saw no evidence of contamination in our samples and most loci present in Black Creek were also present in the Captive Cohort, indicating that there was no bias associated with DNA sampling method in our study. Furthermore, there were no detectable differences in the quality or quantity of sequences obtained from the two sample collection methods (swab and tissue clips). The STACKS pipeline identified 1,185,792 SNPs across 20,464 loci, further filtered by *populations* program constraints to 2,563 putatively independent variant SNPs across the wild populations. When multiple SNPs per locus were allowed, the number of variant SNPs recovered across all wild populations increased to 7,601. After initial runs of the STACKS pipeline, four individuals having less than half the identified SNPs found in the rest of the samples (Jordan18, Captive17, Loving20, and Brown5) were removed, resulting in datasets generated for downstream analyses which included 90 individuals from the original set.

The vast majority of markers sampled were not variable across the four wild populations; out of ~20,000 potential markers (i.e. SNPs), about 10%, met the population parameters specified and contained informative variation. Average nucleotide diversity (π) ranged from 0.21-0.23. Observed heterozygosity was considerably lower than expected heterozygosity across all sites (H_obs_ = 0.08-0.09, H_exp_ = 0.20-0.22), likely indicating genetic bottlenecks at all sites. Allelic richness was also similar across all sampled groups (1.16-1.51). Analysis of the captive population generated 2,416 variable SNPs (8,069 when multiple SNPs per locus were allowed) with an average observed H_obs_ of 0.10 and an average nucleotide diversity of 0.25. Pairwise F_ST_ values were low, with the Brown Creek population presenting the highest differentiation from other populations (0.063-0.082; see Table 1 for all pairwise F_ST_ values); the captive cohort and the Black Creek population had the lowest pairwise F_ST_ (0.035). F_IS_ values for each collection site ranged from 0.44 to 0.48 and was 0.39 for the captive cohort.

Genetic structure was seen among collection sites to varying degrees depending on analytical method. AMOVA was significant at all hierarchical levels (*p* = 0.001), but only 3.89% of genetic variation was explained by whether the sampling site was north or south of the Red River. A further 6.7% of genetic variation was explained by collection sites within regions, indicating landscape barriers to gene flow that are more complex than one major river. The Mantel test and multiple regression for isolation-by-distance was significant (*p* < 0.05), suggesting isolation by distance patterns in the data. DAPC analysis of wild individuals indicated that a K of 4 most accurately captured the diversity of the samples, with individuals from Jordan and Black (Grant Parish) overlapping and Brown and Loving (Rapides Parish) each forming their own distinct cluster (Fig. 4a). When all populations were included, captive mussels overlapped entirely with the Black Creek cluster (Fig. 4b). Notably, the captive mussels showed similar spread in the DAPC analysis as wild individuals from Black Creek, indicating the captive cohort possesses variation in genetic diversity comparable to wild populations.

**Fig. 4.**
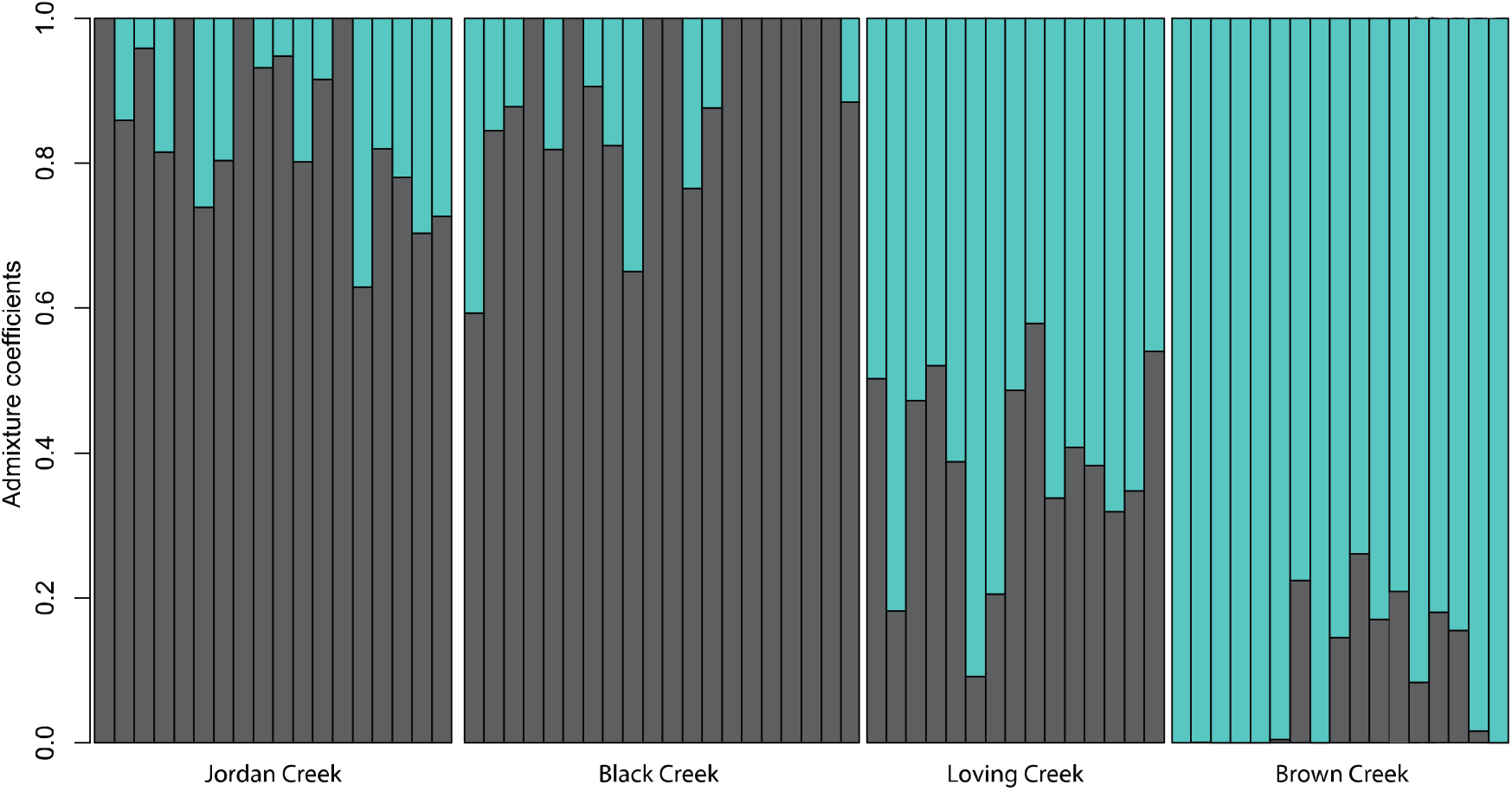
Individual admixtures for all wild individuals as inferred by LEA analysis, using the best-fit K of 2. Colors correspond to parish designations depicted in Fig. 1

Analysis with LEA suggested data were best explained by two genetic clusters when only wild populations were included. LEA analysis with K = 2 showed that individuals from sites north of the Red River (Grant Parish) had more similar admixture profiles to each other than those from south of the Red River (Rapides Parish) and *vice versa* (Fig. 5). Hierarchical clustering analysis with fineRADstructure, mirrored DAPC and LEA analyses (Fig. 6). Brown Creek and Loving Creek formed groupings, to the exclusion of most other individuals, with particularly high coancestry values detected within the Brown Creek population. Individuals from Jordan Creek, Black Creek, and the Captive cohort are nearly indistinguishable from each other with only minor internal structuring comprising a subset of Black Creek and Captive individuals (Fig. 6). When examined in isolation, the captive cohort displayed structured relationships at varying levels of coancestry; at the top of the coancestry value range a cluster of individuals (Cap3, Cap10, and Cap4) may represent a grouping of full siblings (Fig. 7).

**Fig. 5.**
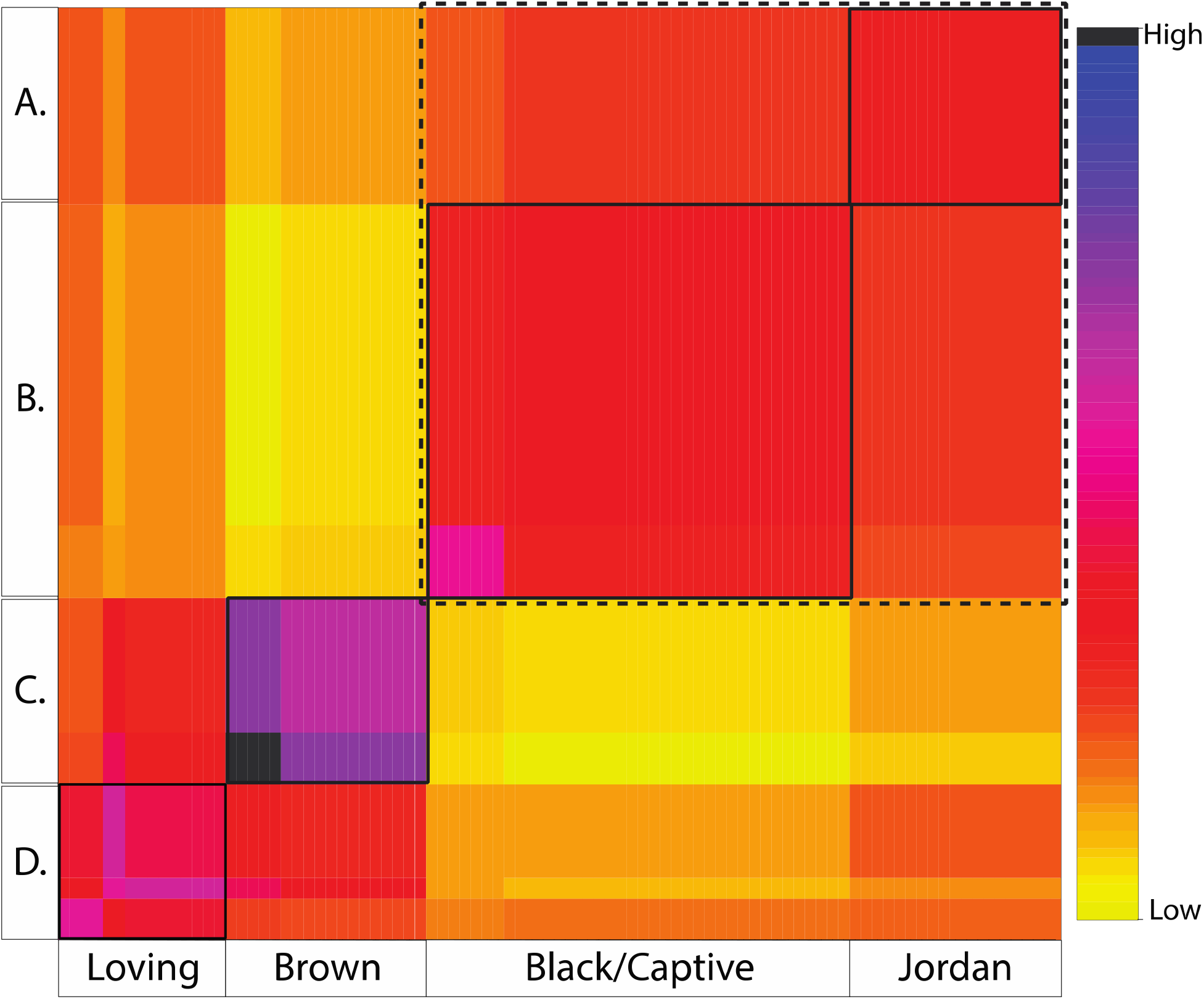
Hierarchical heatmap generated by fineRADstructure, all individuals included. Letters correspond to population identifiers (A = Jordan Creek, B = Black Creek/Captive cohort, C = Brown Creek, D = Loving Creek). Boxes highlight patterns of population clustering, colors represent relative co-ancestry values averaged by population.

**Fig. 6.**
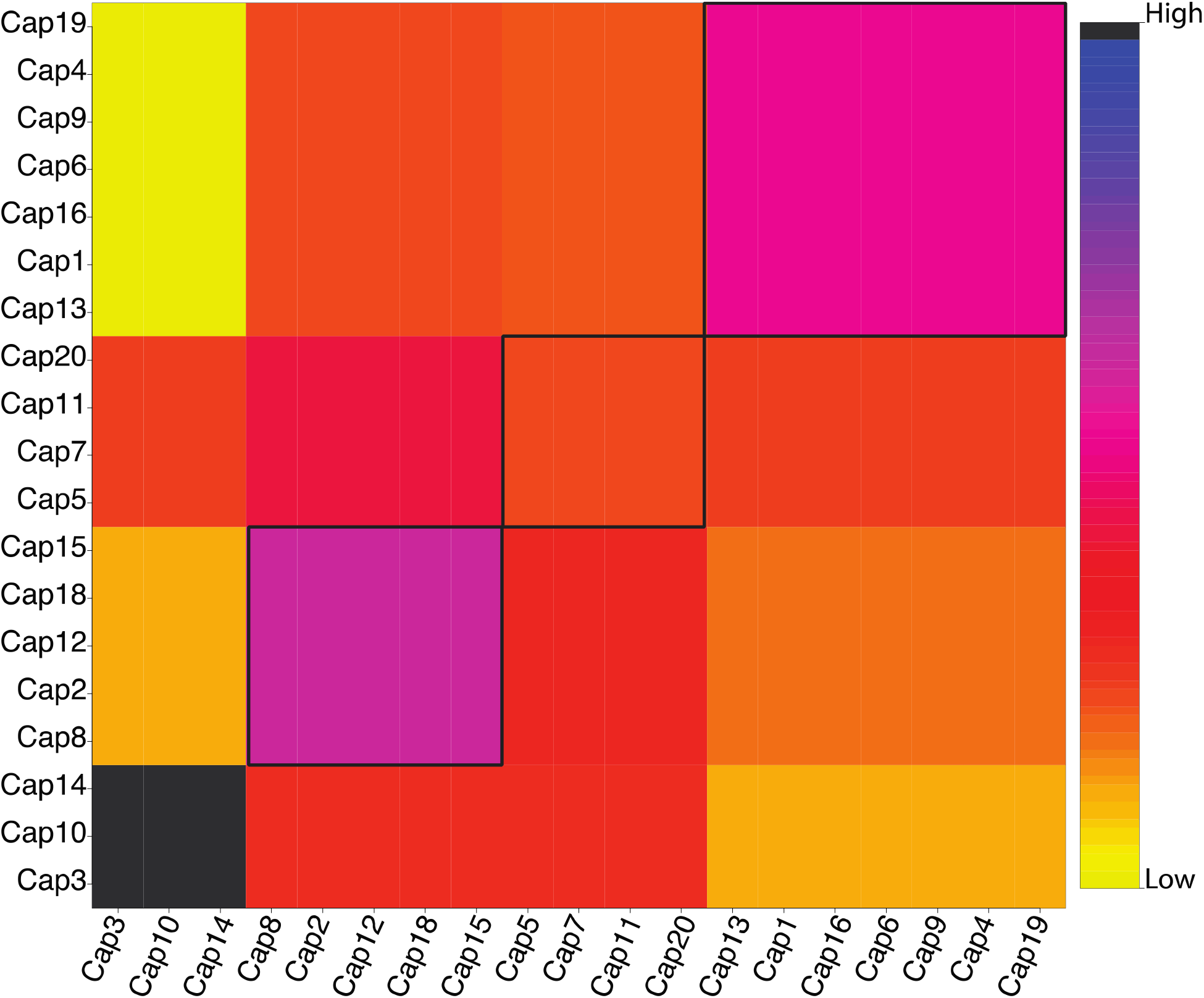
Hierarchical heatmap generated by fineRADstructure, only captive individuals. Boxes highlight patterns of individual clustering, colors represent relative co-ancestry values averaged by cluster.

**Fig. 7.**
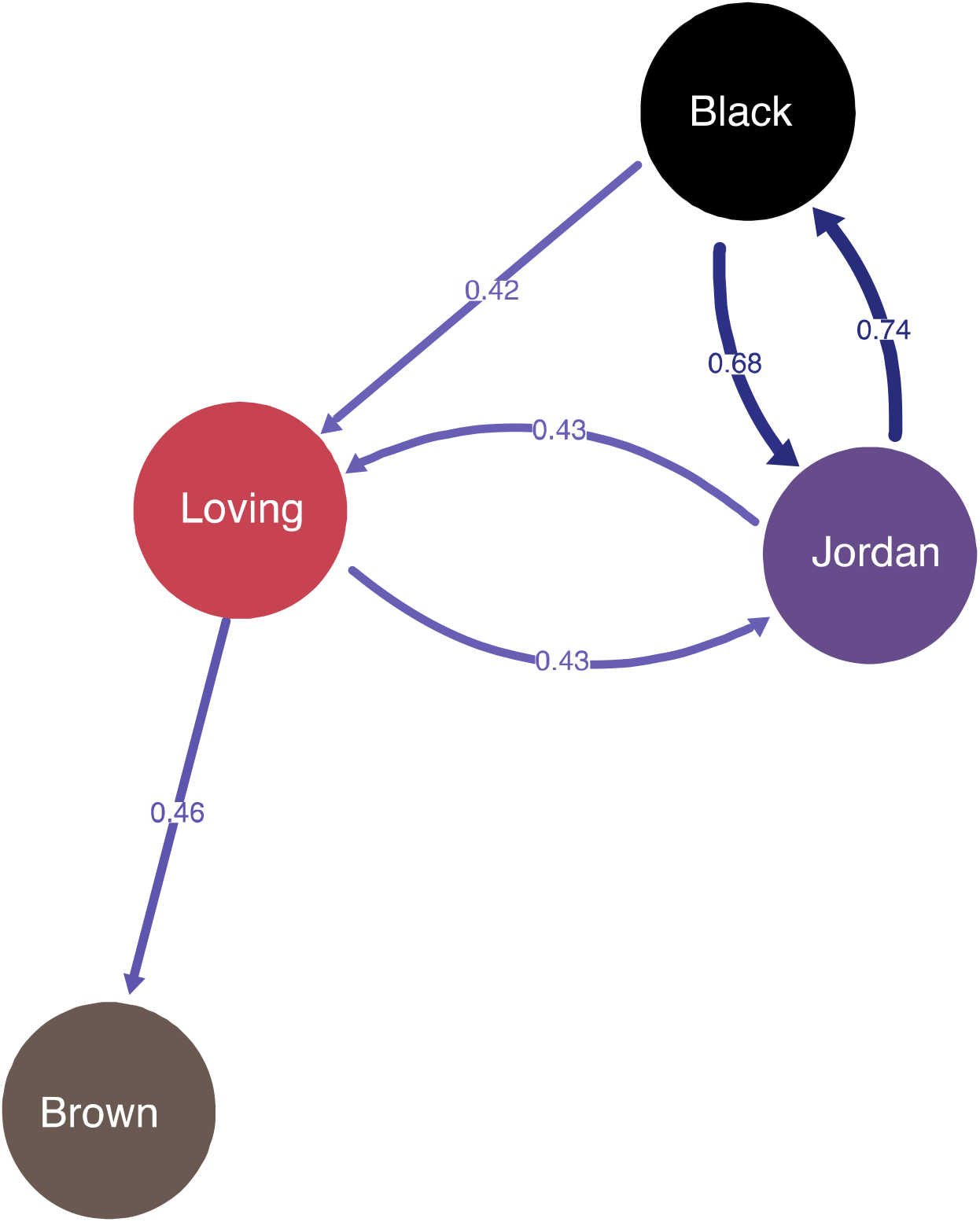
Network generated by divMigrateOnline using the WithCaptiveS dataset; GST statistic, 0.40 filtering level, alpha 0.05. Colors correspond to those in Fig. 3.

The program *divMigrateOnline* detected relatively high migration rates between the two northern sampling sites, Jordan Creek and Black Creek. To a lesser degree, the analysis showed migration from the northern populations to Loving Creek and unidirectional movement from Loving Creek to Brown Creek (Fig. 8). The asymmetry of migration rates inferred by *divMigrateOnline* were not supported by 1000 bootstrap replicates, meaning that while gene flow was detected between Jordan Creek and Black Creek, no strong directionality could be inferred. Of the models assessed by migrate-n, the most likely was model 3 (Table 3) with populations occurring north of the Red River being panmictic, bidirectional gene flow from the panmictic northern population to Loving Creek, and bidirectional migration between Loving Creek and Brown Creek.

Estimates of effective population size for some populations in isolation resulted in negative or infinite values, suggesting that the values were driven by sampling error as a result of insufficient sample size or marker informativeness (Marandel et al. 2019). Combining individuals from the northern sites (Jordan Creek, Black Creek) and southern sites (Brown Creek and Loving Creek) allowed for more realistic estimation of effective population sizes (Table 2). The captive cohort generated the lowest estimates of N_e_ (86 −111 individuals) but an N_b_ of 5. This high estimate for the number of effective breeders within a cohort known to originate from a single female is a strong indication of multiple paternity.

**Table 2.** Population summary statistics (in part, full summary available online) from left to right; number of individuals (N), number of loci recovered (Loci), number of private alleles (PA), nucleotide diversity (Π) and standard deviation (SD), observed heterozygosity (H_o_), expected heterozygosity (H_e_), coefficient of inbreeding (F_IS_), effective populaiton size (N_e_) and 95% confidence interval (CI), number of effective breeders (N_b_), and allelic richness (AR).

| Population | N | Loci | PA | $\Pi$ (SD) | $H_o$ (SD) | $H_e$ (SD) | $F_{IS}$ (SD) | $N_e$ (CI) | $N_b$ | AR (CI) |
| --- | --- | --- | --- | --- | --- | --- | --- | --- | --- | --- |
| **Jordan** | 18 | 2134 | 91 | 0.25<br>($\pm 0.04$) | 0.10<br>($\pm 0.02$) | 0.24<br>($\pm 0.03$) | 0.45<br>($\pm 0.17$) | INF | 4.2 | 1.47 (1.41-1.51) |
| **Black** | 20 | 2203 | 88 | 0.25<br>($\pm 0.04$) | 0.10<br>($\pm 0.02$) | 0.24<br>($\pm 0.04$) | 0.43<br>($\pm 0.16$) | 205 (188-225) | 3.9 | 1.51 (1.46-1.55) |
| **North** | 38 | -- | -- | -- | -- | -- | -- | 177 (170-183) | 3.9 | -- |
| **Loving** | 16 | 1672 | 40 | 0.27<br>($\pm 0.04$) | 0.09<br>($\pm 0.02$) | 0.25<br>($\pm 0.04$) | 0.46<br>($\pm 0.17$) | INF | 4.3 | 1.16 (1.12-1.19) |
| **Brown** | 17 | 1941 | 142 | 0.27<br>($\pm 0.04$) | 0.09<br>($\pm 0.01$) | 0.26<br>($\pm 0.03$) | 0.49<br>($\pm 0.17$) | INF | 3.9 | 1.35 (1.30-1.38) |
| **South** | 33 | -- | -- | -- | -- | -- | -- | 488 (429-566) | 4.1 | -- |
| **Captive** | 19 | 2024 | 41 | 0.25<br>($\pm 0.04$) | 0.10<br>($\pm 0.02$) | 0.24<br>($\pm 0.04$) | 0.40<br>($\pm 0.16$) | 70 (64-78) | 5.0 | 1.37 (1.33-1.41) |

**Table 3.** Description of models assessed by migrate-n, sorted by model rank; model numbers correspond to those in Fig. 2.

| Number | Model Description | Log marginal likelihood value | Rank |
| --- | --- | --- | --- |
| **3** | **Northern panmixia, bidirectional South** | **-93595.56** | **1** |
| 4 | Northern panmixia, full migration | -93803.10 | 2 |
| 1 | Full migration | -94306.55 | 3 |
| 5 | Panmixia | -94030.00 | 4 |
| 2 | Northern panmixia, unidirectional South | -94612.37 | 5 |

COLONY also indicated multiple paternity was present in the captive cohort samples, but the point estimates of male input varied with the level of missing data allowed into the analysis. Using the most complete dataset (sites every individual shared), COLONY output indicated several full-sibling clusters and 7 fathers. The most relaxed filtering strategy (50% site coverage) indicated a different father for each individual in the sample (n = 19). Estimates of N_e_ produced by COLONY (N_e_ = 4, 95% CI = 2-12) were stable across analyses and reflected both the NeEstimator2 values for the captive samples (Table 2) and the fineRADstructure clustering analysis (Fig. 7).

## Discussion

Population genomic data revealed low levels of heterozygosity across *M. hembeli* populations and complex patterns of gene flow among sites. Although low levels of genetic diversity in wild populations is concerning from a conservation standpoint, our results are similar to past studies on *M. hembeli* (Curole et al. 2004; Roe 2009). Moreover, low genetic diversity and a signature of bottlenecks at every sampling site may reflect natural processes such as rapid colonization after the loss of a habitat that results from stream meanders and cutoffs that are common in low-elevation, flat terrains like those in central Louisiana. Despite low genetic diversity, high gene flow among some sample sites was observed. Genetic structure generally followed a pattern of isolation-by-distance, but the Red River also appeared to be a factor in observed genetic structure. This suggests movement between populations on the same side of the Red River may occur during major flood events, thereby facilitating migration with infected fishes moving to adjacent stream channels and does not require host-fish passage through the Red River channel.

Our data showed a captive cohort of *M. hembeli* produced from a single female had comparable genetic diversity to the wild population from which the female was sampled. Broadly, analyses indicate evidence of multiple paternity, which has considerable implications for propagation efforts of *M. hembeli*. Although multiple paternity has been known to occur in at least some freshwater mussels (Christian et al. 2007; Bai et al. 2012; Ferguson et al. 2013; Wacker et al. 2018), more recent studies have empirically demonstrated wild female mussels can mate with multiple males to produce genetically diverse captive cohorts (Wacker et al. 2019). Multiple paternity had not been empirically observed in *M. hembeli* prior to this study. Results suggest progeny generated from a few (randomly selected) wild-fertilized females annually would mirror the heterogenity found in the wild host populaton, likely resulting in low detectable differences between propagated and wild mussels with a recovery effort that spans multiple years.

### Genetic Diversity and Effective Population Size

Low observed heterozygosity relative to expected heterozygosity (Table 2), suggests *M. hembeli* populations are small and experiencing associated effects such as inbreeding and genetic bottlenecks. This is also reflected by high F_IS_ values (Table 2). Given the threatened status of *M. hembeli* under the U.S. Endangered Species Act, anthropogenic activities have clearly caused severe population declines. However, life history, natural demographics, and stochastic habitat disturbances probably also play a possibly an outsized role, in influencing low heterozygosity of *M. hembeli*. For instance, a single beaver dam once led to the extirpation of an *M. hembeli* bed (~1000 individuals) located on Forest Service land (Stewart 1990). Furthermore, Johnson and Brown (2000) showed that channel stability of the study area on the time-scale of a single year can be low, even though they noted that *M. hembeli* appeared to be associated with relatively stable microhabitats. Assuming rapid colonization ability of *M. hembeli* (see Curole, Foltz, and Brown 2004), low heterozygosity is likely the result of repeated bottlenecks caused by habitat destruction followed by colonization of newly exposed suitable habitat. Although natural destruction of mussel beds and rapid colonization of new habitats may be a normal part of *M. hembeli* biology, habitat fragmentation caused by anthropogenic activity exacerbates contemporary population decline by restricting recruitment across populations (Geist and Auerswald 2007). The habitat fragmentation witnessed through modification of waterways or changes in landuse are likely limiting re-colonization options for *M. hembeli*.

When considered in isolation, N_e_ estimates for all sampling sites except Black Creek were inferred to be infinite, which is indicative of N_e_ estimates being driven by sampling error, rather than large population sizes (Marandel et al. 2019). Pooling individuals from north and south of the Red River, respectively, allowed for more precise estimates of N_e_ (Table 2). Effective population size is difficult to accurately estimate even with genome-wide markers as many methods make assumptions that are typically violated such as sampling of non-overlapping generations (Waples et al. 2016; Wang et al. 2016; Marandel et al. 2019). The linkage disequilibrium method applied here is known to be downwardly biased (as much as 30%) when samples consist of different age classes (Waples et al. 2014). We anticipate that our estimates of N_e_ for wild populations are much lower than reality, as multiple age classes were sampled. Thus, N_e_ estimates reported here may be of value to resource managers, but they should be approached with caution and not the sole genetically derived metric used for assessing populations of *M. hembeli*. The temporal trajectory of N_e_ is more crucial to conservation managers than a point estimate, and we recommend additional sampling of size/age class cohorts to reveal trends through time.

### Population Connectivity

In general, analyses determined genetic demarcation between populations occurring north of the Red River in those occurring south of it. However, fine-scale relationships and migration patterns among populations that our SNP-based approach illuminated are novel. Sampled sites appeared to demonstrate an isolation-by-distance effect, which is a general pattern seen in many freshwater organisms (Meffe and Vrijenhoek 1988; Whelan et al. 2019). Furthermore, AMOVA indicated significant genetic structure with a genetic break occurring between collection sites north and south of the Red River, with further significant genetic structure between populations in the north and south. These patterns were recovered to varying degrees by clustering analyses, with DAPC indicating at least some genetic distinctness among all four sites (Fig. 4).

Analyses that examined finer-scale gene-flow patterns indicated a high amount connectivity between Grant Parish populations (Jordan Creek and Black Creek), with the best-fit model as assessed by migrate-n having a panmictic Jordan Creek and Black Creek (Fig. 2).Whether the observed pattern is a result of active gene flow or a historical connection is unknown. Black Creek has been putatively isolated from Jordan Creek for the past 60 years behind Iatt Lake, but the long lifespan of *M. hembeli* may result in a longer time-period for genomic signatures of isolation to be detectable. Documentation of Iatt Lake’s management history (Moses et al. 2016) indicates *Esox americanus americanus*, was not detected in fish surveys but other *Esox* species (*Esox americanus vermiculitus*, *Esox niger*) have been collected during Iatt Lake surveys. Iatt Lake is prone to flooding and has experienced several high water events in recent history (Moses et al. 2016), which could facilitate fish passage between tributaries. Assuming the presence of *Esox* spp. around Lake Iatt and periodic flooding it is possible that a connection was recent and possibly intermittent between Black Creek and Jordan Creek drainages.

Relative genetic homogeneity among northern sites contrasts with the relative isolation of those sampled from south of the Red River. Loving Creek and Brown Creek shared similar genomic backgrounds, as indicated by LEA (Fig. 4), but *divMigrate* and migrate-n analyses indicated somewhat limited gene flow between the two southern sites. Several analyses indicated that Brown Creek was the most isolated group of individuals sampled, which may be partially explained by a higher stream distance from its Red River confluence compared to other sites. Connectivity analyses appear to indicate that Loving Creek represents Brown Creek’s only connection to the rest of the species range (Fig. 6). Though there are sites not sampled during the course of this study which occur in streams located between Loving and Brown Creek, our data still indicates that movement of *M. hembeli* in the southern part of its range is relatively more restricted than in the northern section of its range.

Further natural history work is needed to better understand the conditions required for successful dispersal of *M. hembeli*. Although a detailed host fish trial has not been completed for *M. hembeli*, formal trials have been completed for *M. marrianae*, the Alabama Pearlshell. These trials indicate *Esox vermicularis vermicularis* is the primary host in Alabama. A minimal transformation rate was also documented for *Noturus leptacanthus* (Speckled Madtom) but it’s not a primary host for *M. marrianae* (Fobian et al., *in prep.*). In contrast, *M. hembeli* readily transforms transform on *Esox* spp. in captivity (Schmidt-Frater, *pers. comm.*) but a formal host trial has not been completed. Additionally, several non-sampled populations of *M. hembeli* occur in headwater streams isolated behind reservoirs – future studies including those sites will likely provide further insights into the impact of such barriers on the dispersal of *M. hembeli* across its range.

### Captive Propagation and Reintroduction

Our analysis of captively reared individuals revealed a surprising amount of genetic diversity given that the cohort was produced by a single female from Black Creek. Notably, genetic diversity estimates for this cohort were virtually identical to estimates from the wild population (Table 2). Clustering analyses provided evidence these data were more than sufficient to assign captive individuals to their population of origin, always grouping them with wild Black Creek individuals (Fig. 4b). Multiple paternity was also evident based on estimates of the number of effective breeders for the captive cohort (N_b_ =5). Coupled with inferences from COLONY and fineRADstructure, where multiple paternal genotypes (7-19) and multiple clusters of individuals with high (but not identical) co-ancestry were observed, our study supports the presence of a multiple paternity strategy within *M. hembeli*. Overall, this represents a best-case scenario for managers as a limited number of wild-caught *M. hembeli* females can be brought into captivity annually and produce a genetically diverse cohort for reintroduction efforts. However, given the difficulty of *Margaritifera* spp. production in a hatchery setting (Paul Johnson *pers. obs.*), any serious reintroduction effort would likely be a decades-long endeavor at minimum.

The ability to produce genetically diverse individuals from a small number of females should facilitate propagation programs; however, care must still be taken when choosing brood stock and determining reintroduction sites. Managers should be guided by our findings of population structure and isolation by distance. Broadly, brood stock should be as geographically proximate to the chosen reintroduction site as possible, coming from one or a few sites within the same drainage. At the very least, broodstock should come from the same side of the Red River as the chosen reintroduction site. Furthermore, genetic diversity and number of effective breeders for another Margaritiferid, *M. margaritifera*, was recently determined to be higher when females were fertilized in the wild relative to those fertilized in captivity (Wacker et al. 2019). Our work suggests multiple paternity is likely the case for *M. hembeli* as well. The best chance we have at maintaining appropriate levels of diversity is to utilize wild-fertilized broodstock while it is still available, rather than attempt to establish a captive breeding colony of *M. hembeli*.

Our data also give reasons to avoid augmentation (i.e., placing captively reared individuals on top of a natural population) in favor of reintroductions (i.e., placing captively reared individuals at a site from which *M. hembeli* has been extirpated). Each population analyzed here was considered genetically distinct in at least one analysis, albeit with limited or no genetic distinction between Black Creek and Jordan Creek in most analyses. That said, consequences of outbreeding depression are impossible to predict at this time, and augmentation violates recent recommendations for freshwater mussel propagation and release (Mobile River Basin Mollusk Restoration Committee 2010; Cumberlandian Region Mollusk Restoration Committee 2010; Strayer et al. 2019). Our data support such recommendations. If managers are faced with no suitable sites for propagation and release other than sites with natural *M. hembeli* population, then we argue that habitat restoration should be a higher priority than captive propagation of *M. hembeli*. Good animals placed into poor habitat will not have the desired outcome (Geist and Auerswald 2007).

## Conclusions

This study provides information that can be used to facilitate successful propagation efforts and a framework for studying the existing diversity in imperiled mussel species using modern methods. We have demonstrated that genetically diverse cohorts of margaritiferids may be produced from a small number of wild-caught, gravid females. Importantly, our findings also indicate that the occurrence of multiple paternity in freshwater mussels may be more widespread than the limited number of explicitly documented cases. Our findings can likely be generalized to closely related species such as the federally endangered Alabama Pearlshell, *M. marrianae,* the sister species to *M. hembeli*, which is currently the focus of intense propagation and management efforts. More broadly, we have demonstrated the utility of RAD-seq approaches, compared to older technologies, in providing fine-scale information for freshwater mussel conservation. Although RAD-seq is now widely used for many conservation genetics studies of non-model organisms, its use for freshwater mussel research is still rare.

In terms of Louisiana Pearlshell recovery, our recommendation is to continue propagation efforts utilizing wild-fertilized females with a focus on habitat restoration and continued life history research. Given the likelihood of multiple parentage, captive cohorts produced from a single female will have more diversity than might have been previously expected. However, efforts should be made to not re-use the same females over multiple propagation years, and the use of multiple broodstock females per year is encouraged when possible. Broodstock selected from populations north of the Red River should not be used to propagate individuals that are to be released into localities south of the river. While our findings provide some hope for the efficacy of propagating the Louisiana Pearlshell using existing populations as sources of diverse brood stock, it also indicates that high levels of inbreeding and loss of population connectivity may be a looming problem for long-term survival of the species. Indeed given the genetic bottlenecks at multiple sites sampled in this study, further analyses might reveal hererogeneity of a reintroduction effort could be improved by mixing propagules from multiple adjacent populations, taking care to keep individual efforts within their respecitive subdrainages. More work is needed to ensure that reintroduced and existing populations of this threatened species form a connected, contemporarily recruiting network of individuals capable of sustaining itself if true recovery is to be achieved.

## Acknowledgements

We thank Jared Streeter (Louisiana Department of Wildlife and Fisheries), Ted Soileau, and Steve Shively (US Forest Service) for assistance in the field. Nathan Johnson (US Geological Survey) and Jürgen Geist (Technische Universität München) provided advice on swabbing and extraction methods. A special thanks to the staff of the Alabama Aquatic Biodiversity Center for maintaining captively reared *M. hembeli*, developing propagation techniques for the species, and allowing access to tissues. We also thank the landowners who provided access to sampling sites on privately owned land. The findings and conclusions in this article are those of the authors and do not necessarily represent the views of the U.S. Fish and Wildlife Service.

## Funding

This work was funded by a 1311 system-wide funding grant from U.S. Fish and Wildlife Service.

## Availability of data and material

Demultiplexed Illumina sequence data have been uploaded to NCBI SRA (ascession numbers to be provided upon manuscript acceptance). Processed datasets in various file formats and certain program input and output files (e.g., colony) are available on FigShare (private link for reviewers:; *DOI to be provided upon acceptance*). Additional code related to the execution of pipelines used are available online at https://github.com/nlgarrison/ConservationGenomics.

## Notes

#### Summary of Updates

removed private link, fixed author order

